# The hydrophobic nature of dengue 2 M residue 36 influences E protein and ApoptoM peptide

**DOI:** 10.1101/2023.02.07.527418

**Authors:** Jason Decotter, Philippe Desprès, Gilles Gadea

## Abstract

The recent epidemics of dengue in South West Indian Ocean coincided with the emergence of Cosmopolitan dengue virus type 2, including viral strains DES-14 in Tanzania and then RUN-18 in La Reunion. The initial step of dengue virus assembly is the formation of heterodimers between prM and E proteins where prM acts as a chaperone for E. During dengue virus maturation, prM is cleaved into membrane protein M which embeds a pro-apoptotic peptide consisting of residues 31/41 and referred as ApoptoM. An infrequent valine at position M-36 was found in DES-14 whereas RUN-18 bears a common isoleucine. Here, we investigated whether Ile-to-Val substitution at position M-36 may have an impact on RUN-18 E expression and cell-death promoting capability of RUN-18 ApoptoM. Using recombinant RUN-18 envelope proteins expressed in human epithelial A549 cells, we showed that Ile-to-Val but not Ile-to-Ala substitution affects the behavior E protein reducing the cytotoxicity of prM and E proteins. The substitution of isoleucine by valine at position M-36 leads to increase the apoptosis-inducing activity of ApoptoM. Our data highlight that hydrophobic nature of amino-acid residue in position 36 of dengue virus M protein influences E protein expression and death-promoting activity of ApoptoM opens up important perspectives in the development of effective live-attenuated DENV vaccines.

## INTRODUCTION

Dengue is a major public health problem worldwide, particularly in tropical and subtropical regions [1,2]. Dengue disease is caused by dengue virus (DENV), a mosquito-borne enveloped RNA virus. DENV belongs to the *Flavivirus* genus (*Flaviviridae* family) which includes other medically important arthropod-borne viruses such as Japanese encephalitis virus (JEV), tick-borne encephalitis virus (TBEV), West Nile virus (WNV), yellow fever virus (YFV), and Zika virus (ZIKV). There are four genetically related but antigenically distinct serotypes of DENV named DENV-1, DENV-2, DENV-3 and DENV-4 respectively. DENV infection can result in a wide range of clinical manifestations, from flu-like disease (dengue fever) to severe illness (severe dengue), a potentially lethal disease [3,4]. In 2018-19, Reunion Island has experienced a dengue epidemic with nearly 25,000 reported cases. The predominant isolated serotype was DENV-2, followed by DENV-1 and DENV-3 [5–7]. DENV-2 strains isolated from dengue patients in 2018 (Reunion/2018) were sequenced and phylogenetic analysis reported that all DENV-2 clinical isolates from Reunion/2018 belong to the Cosmopolitan genotype (or genotype II), including the prototypical RUJul strain (referred as RUN-18, GenBank accession number MN272404.1) [8,9]. In the South-West Indian Ocean (SWIO) region, other epidemic DENV-2 strains of cosmopolitan genotype were also sequenced including the epidemic DENV-2 strain D2_K2_RIJ_059/Dar es Salaam 2014 (referred as DES-14, GenBank accession number MG189962.1) isolated from a patient diagnosed for dengue fever in Dar-es-Salaam, Tanzania, in 2014 [10]. DENV contains a positive-sense single-stranded RNA encoding a single large polyprotein precursor that is co- and post-translationally processed by host and viral protease into three structural proteins which are the capsid protein (C), the precursor membrane protein (prM) and the envelope protein (E), followed by seven nonstructural proteins. DENV E glycoprotein (495 amino-acid residues) is involved in virus attachment to host-cell, in internalization of virus particles, and then in the fusion between viral and cellular membranes releasing genomic RNA into the cytosol. The E protein includes an ectodomain followed by a stem region and two transmembrane domains (TMDs). The E ectodomain comprises three domains: a central β-barrel shaped domain I (EDI), a finger-like domain II (EDII), and a C-terminal immunoglobulin-like domain III (EDIII). DENV fusion peptide (residues 98 to 112) has been identified in domain II. DENV prM glycoprotein (166 amino-acid residues) consists of “pr” peptide (residues 1 to 91) followed by the surface structural M protein (75 aminoacid residues). The initial step of virus assembly involves heterodimeric interactions between prM and E proteins, prM playing the role of chaperone for E to prevent a premature fusion of immature virus particles during their transport through the secretory pathway. The heterodimeric interactions between prM and E proteins are important for the proper folding of E. The processing of prM occurs in the trans-Golgi apparatus by cellular furin/furin-like protease family that cleaves the “pr” peptide from the surface membrane M protein [11–13]. DENV M protein consists of a N-terminal ectodomain (residues 1 to 40, hereafter referred as ectoM) which is formed of a flexible hydrophobic loop followed by an amphipathic peri-membrane α-helix and ended by a C-terminal transmembrane region [14]. In the topology of prM-E heterodimer, the M ectodomain appears to be adjacent to EDII domain [14,15]. DENV-2 M protein includes a pro-apoptotic ApoptoM peptide involving the residues M-32/40 [16]. The cell death-inducing activity of ApoptoM involves caspase-3 activation and its transport in the secretory pathway is a prerequisite for initiation of apoptosis [16,17]. Amino-acid residue in position 36 of M protein has been identified as playing an important role in flavivirus replication cycle [18–20]. It has been reported that M-36 amino-acid substitutions affect virus assembly and/or egress and thus, the pathogenicity of flaviviruses [19,20]. Moreover, the M residue 36 influences the cell death-promoting capability of ApoptoM [16]. Comparative sequence analysis between the M proteins of DENV-2 strains RUN-18 and DES-14 identified a single amino-acid substitution at position M-36. Interestingly, an infrequent valine residue has been identified at position M-36 in DES-14 whereas RUN-18 has a common isoleucine. In the present study, we investigated the effects of Ile-to-Val substitution at position M-36 on RUN-18 E folding and pro-apoptotic ApoptoM peptide using recombinant viral proteins were expressed in human epithelial A549 cells.

## MATERIALS and METHODS

### Cell Culture, Antibodies and Reagents

For cell culture, human epithelial A549 cells (ATCC, CCL-185) were cultured in Dulbecco’s Modified Eagle’s Medium (DMEM) supplemented with 10 % heat-inactivated fetal bovine serum (FBS) (Dutscher, Brumath, France), 2 mmol.L^−1^ L-Glutamine, 1 mmol.L^−1^ sodium pyruvate, 100 U.mL^−1^ of penicillin, 0.1 mg.mL^−1^ streptomycin and 0.5 ug.mL^−1^ of fungizone (Amphotericin B) (PAN Biotech, Aidenbach, Germany) at 37 °C under 5 % CO_2_ atmosphere. The mouse anti-pan flavivirus envelope E glycoprotein domain II (EDII) mAb 4G2, purchased from RD Biotech (Besançon, France), and the mouse anti-DENV envelope E glycoprotein domain III (EDIII) mAb 4E11, a kind gift of Pasteur Institute (Paris, France), were used for immunodetection of the viral envelope E protein. Mouse anti-6x(His) mAb was purchased from Abcam (Cambridge, UK), to detect NS1 protein expression. Rabbit anti-calnexin pAb was purchased from Santa-Cruz Biotechnology (Clinisciences, Nanterre, France) and rabbit anti-β actin pAb from ABclonal (Massachusetts, USA), to detect the ubiquitously and constitutively expressed Calnexin and β actin proteins respectively. Donkey anti-mouse Alexa Fluor 488 and donkey anti-rabbit Alexa Fluor 594 secondary antibodies were purchased from Invitrogen (Carlsbag, CA, USA). Goat anti-mouse and goat anti-rabbit immunoglobulin-horseradish peroxidase (HRP) conjugated antibodies were purchased from Abcam (Cambridge, UK). Blue-fluorescent DNA stain 4’,6-diamidino-2-phenylindole (DAPI) was purchased from Euromedex (Souffelweyersheim, France). Lipofectamine 3000 (Thermo Fisher Scientific, Les Ulis, France) was used for transfection, according to the manufacturer’s instructions.

### Vector Plasmids Expressing Recombinant Dengue proteins

A mammalian codon-optimized synthetic gene coding for the RUN-18 prM and E proteins as a polyprotein (Genbank accession number MN272404.1) and preceded by the RUN-18 prM signal peptide was chemically synthetized and then inserted into *Nhe* I and *Not* I restriction sites of vector plasmid pcDNA-3.1 by Genecust (Boynes, France). Site-directed mutagenesis was performed on resulting plasmid pcDNA3/RUN-18 prME to generate mutants bearing M-(33A, 35A, 38A), M-(I36A) or M-(I36V) mutation by Genecust (Boynes, France). A mammalian codon-optimized synthetic gene coding for the reporter gene GFP followed by the RUN-18 ectoM (residues M-1/41) or ApoptoM sequence (residues M-31/41) (Genbank accession number MN272404.1) and preceded by the RUN-18 prM signal peptide was inserted into *Nhe* I and *Not* I restriction sites of vector plasmid pcDNA-3.1 by Genecust (Boynes, France). As short Gly-Ser spacer was inserted between sGFP and viral sequences.Site-directed mutagenesis was performed on resulting plasmids pcDNA3/sGFP-ectoM and pcDNA3/sGFP-ApoptoM to generate mutants bearing M-(33A, 35A, 38A), M-(I36A) or M-(I36V) mutation by Genecust (Boynes, France). The production of endotoxin-free plasmids, their quantification, and the sequencing were performed by Genecust (Boynes, France). A549 cells were transiently transfected with plasmids using Lipofectamine 3000 (Thermo Fisher Scientific, Les Ulis, France) according to the manufacturer’s instructions.

### Western Blot Assay

Cells were seeded in 24-well plates at a density of 1.5 × 10^5^ cells per well, then transfected with vector plasmid constructs. After 18 h of transfection, cells lysates were performed in radioimmunoprecipitation assay (RIPA) lysis buffer (Sigma-aldrich Humeau, La Chapelle-Sur-Erdre, France) at 4 °C for 10 min. Lysates were centrifuged 12000 g for 20 min at 4 °C. Proteins from total cells extracts were quantified using bicinchoninic acid (BCA) reagent according to manufactured instructions (Sigma-aldrich Humeau, La Chapelle-Sur-Erdre, France). Protein extracts were treated with Leammli buffer, either under reducing (heated at 95 °C for 5 min, in presence of dithiothreitol (DTT)) or nonreducing conditions (no heat, no DTT). Proteins were separated by in house 4-12 % SDS-PAGE gel and transferred onto nitrocellulose NC Protan membrane with a 0.45 μM pores (Amersham, GE, Buc, France). The membranes were blocked for 20 min with 5 % non-fat dry milk in PBS-Tween, and incubated with anti-4G2, anti-6x(His) or anti-β actin antibodies (dilution 1:2000) for 1h at room temperature. Anti-mouse or anti-rabbit IgG HRP-conjugated secondary antibodies were used at 1:10000 dilution. Membranes were incubated with enhanced chemiluminescence (ECL) prime detection reagents (GE, Buc, France) and exposed using Amersham imager 680 (GE, Buc, Healthcare).

### Flow cytometry Analysis

Cells were seeded in 24-well plates at a density of 1.5 × 10^5^ cells per well, then transfected with vector plasmid constructs. After 18 h of transfection, cells were gently harvested by trypsinization, fixed with 3.7% paraformaldehyde (PFA) in phosphate buffered saline (PBS) for 10 min and then observed for GFP. For detection of the E protein, cells were permeabilized with 0.15 % Triton X-100 in PBS for 5 min, and then blocked with 2 % bovine serum albumin (BSA) in PBS, and then labeled with anti-4G2 or anti-4E11 antibodies (dilution 1:2000) for 1 h at room temperature. For the detection of NS1 protein, cells were labeled with anti-6x(His) antibody (dilution 1:2000). Goat anti-mouse Alexa Fluor 488 IgG antibody was used as a secondary antibody (1:2000) for 30 min. For each assay, 10^4^ cells were analyzed by flow cytometry using a CytoFLEX flow cytometer (Beckman, Coulter, Brea, CA, USA) using CytExpert software (version 2.1.0.92, Beckman Coulter, Villepinte, France).

### RT-qPCR assays

Cells were seeded in 24-well plates at a density of 1.5 × 10^5^ cells per well, then transfected with vector plasmid constructs. Quantification of prM and E RNA was performed by RT-qPCR. After 18 h of transfection, total cellular RNA was extracted from cells with RNeasy kit (Qiagen, Hilden, Germany) according to the manufacturer’s recommendations. Total cDNA was obtained by reverse transcription using random hexamers pd(N)6 and M-MLV reverse transcriptase (Life Technologies, Carlsbad, CA, USA) at 42 °C for 50min. cDNA were amplified using 0.2 μM of each primer (prM: forward primer 5’-CAACAGYGATGGCGTTC-3’, reverse primer 5’-TCCARTCCCATTCCCAC-3’; E: forward primer F5’-GTTACACCCCACTCTGGC-3’, reverse primer 5’-GAAGTCCAGGCCAGTCCG-3’), 2X Absolute Blue qPCR SYBR Green Low ROX Master Mix (ThermoFisher, Waltham, MA, USA) on a CFX96 Real-Time PCR Detection System (Bio-Rad, Life Science, Hercules, CA, USA). The CFX96 software (Bio-Rad) was used to evaluate the threshold cycle (Ct) for each well in the exponential phase of amplification. Results were normalized to the housekeeping gene GAPDH in each condition.

### Immunofluorescence Confocal Assay

Cells were seeded in 24-well plates at a density of 1.5 × 10^5^ cells per well, grown as a monolayer on glass coverslip and were then transfected with vector plasmid constructs. After 18 h of transfection, cells were fixed on coverslip with 3.7% PFA in PBS for 15 min. Cells were permeabilized with 0.15% Triton X-100 in PBS for 15 min, blocked with 2 % BSA in PBS, and then labeled with anti-4G2 and anti-calnexin antibodies (dilution 1: 1000) for 1 h at room temperature. Donkey anti-mouse Alexa Fluor 488 and donkey anti-rabbit Alexa Fluor 594 were used as secondary antibodies (1:2000) for 30 min, and the nucleui were stained with DAPI. 50 % glycerol in PBS was used as mounting media. Capture of the fluorescent signal was allowed with a Nikon Eclipse TI2-S-HU (confocal microscopy) coupled to the imaging software NIS-Element AR (Nikon, Champigny-sur-Marne, France).

### LDH Assay

Cells were seeded in a 96-well culture plate at a density of 2.5 × 10^4^ cells per well. Cytotoxicity was evaluated by quantification of lactate dehydrogenase (LDH) release in culture supernatant of transfected-cells using CytoTox96 nonradioactive cytotoxicity assay (Promega, Charbonnières-les-Bains, France) according to the manufacturer’s instructions. The absorbance of converted dye was measured at 490 nm with background subtraction at 690 nm using a microplate reader Tecan (Tecan Trading AG, Männedorf, Switzerland).

### MTT Assay

Cells were seeded in a 96-well culture plate at a density of 2.5 × 10^4^ cells per well. Cell monolayers were rinsed with PBS and incubated with supplemented DMEM culture growth medium mixed with 5 mg.mL^−1^ MTT (3-[4,5-dimethylthiazol-2-yl]-2,5-diphenyltetrazolium bromide) solution for 1 h at 37 °C. MTT medium was removed, and the formazan crystals were solubilized with dimethyl sulfoxide (DMSO). Absorbance was measured at 570 nm with background subtraction at 690 nm, using a microplate reader Tecan (Tecan Trading AG, Männedorf, Switzerland).

### Caspase 3/7 Enzymatic Activity

Cells were seeded in a 96-well culture plate at a density of 2.5 × 10^4^ cells per well. Caspase 3/7 enzymatic activity in raw cell lysates was measured using a Caspase Glo 3/7 assay kit (Promega) according to the manufacturer’s recommendations. Caspase activity was quantified by luminescence using a FLUOstar Omega Microplate Reader (BMG Labtech, Champigny-sur-Marne, France).

### Statistical Analysis

Our results were statistically analyzed using the GraphPad Prism software (version 9, GraphPad software, San Diego, CA, USA). One-way ANOVA tests were performed to compare quantitative data between the different experimental conditions, with Dunnett or Tukey correction for multiple comparisons. Values of *p* < 0.05 were considered statistically significant for ANOVA test. The degrees of significance are indicated on the figure, **** *p* < 0.0001, *** *p* < 0.001, ** *p* < 0.01, * *p* < 0.05 or ns: not significant. All values were expressed as mean ± SD of at least three independent experiments.

## RESULTS

### Characterization of recombinant RUN-18 E protein co-expressed with prM in A549 cells

As proper folding of DENV-2 E protein involves the chaperone activity of prM [12,14], recombinant RUN-18 E protein was synthesized with prM as a polyprotein precursor which is then processed by cellular signalases. A synthetic gene encoding RUN-18 prM and E with optimized codons for expression in human cells was inserted into the vector plasmid pcDNA3. In the resulting plasmid pcDNA3/RUN-18 prME, the sequence coding for prM followed by E was preceded by the authentic RUN-18 prM signal sequence.

Expression of recombinant RUN-18 E protein was assessed in A549 cells transfected with pcDNA3/RUN-18 prME for 18 h. A plasmid pcDNA3/RUN-18 NS1 encoding recombinant soluble NS1 protein from RUN-18 was used as a plasmid control since protein expression results in a weak effect on cell viability [21]. Expression of recombinant RUN-18 NS1 in A549 cells transfected with pcDNA3/RUN-18 NS1 was verified by FACS analysis and immunoblot assay (Figure S1). The antigenic reactivity of recombinant E protein was assessed by FACS analysis using anti-DENV E monoclonal antibodies (mAbs) 4G2 and 4E11 that bind to epitopes into DENV-2 EDII and EDIII, respectively (Figure 1A). Immunoblot assay using mAb 4G2 allowed the detection of recombinant E protein expressed in A549 cells (Figure 1B).

**Figure 1.**
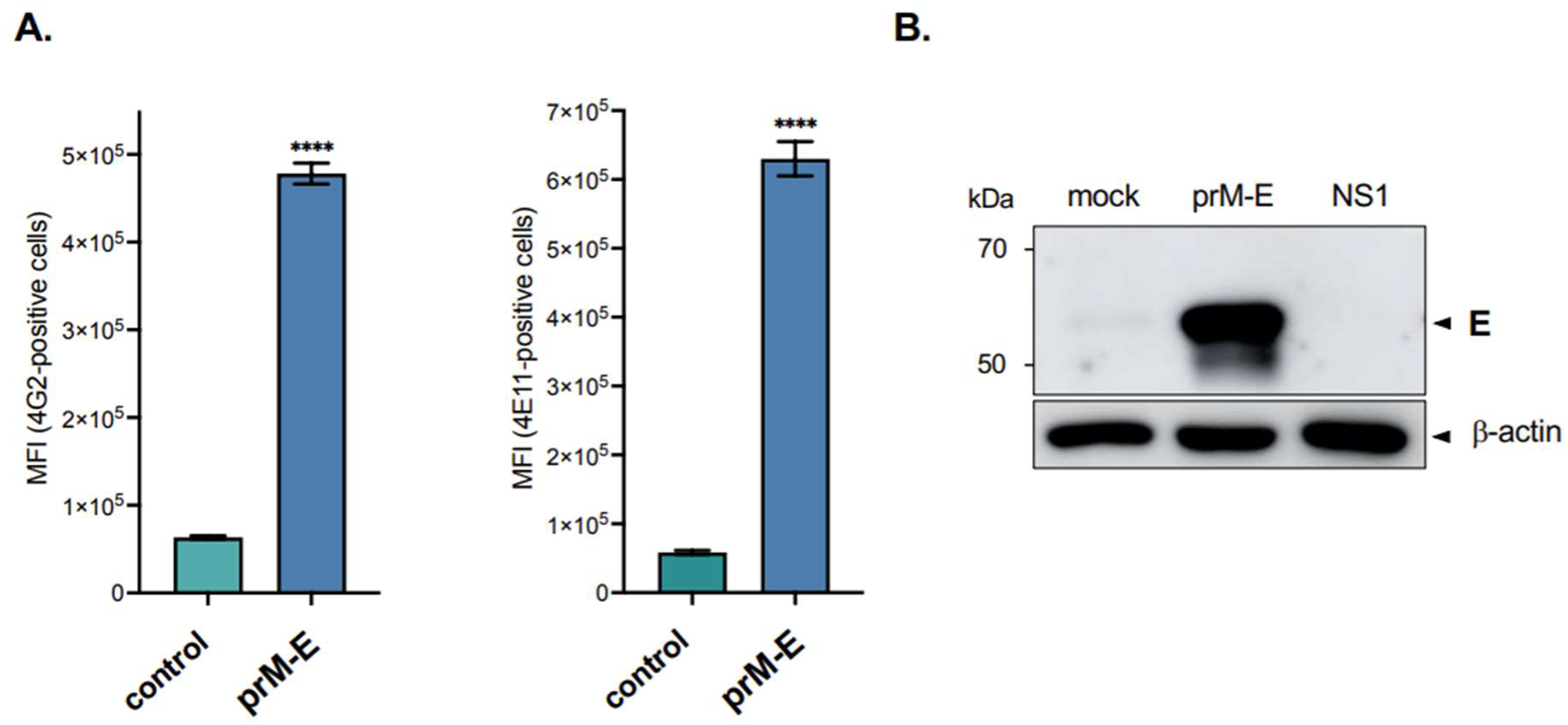
Detection of RUN-18 E protein co-expressed with prM in A549 cells. A549 cells were transfected for 18 h with plasmid pcDNA3/RUN-18 prME expressing recombinant RUN-18 prM and E proteins (prM-E) or mock-transfected (mock). A plasmid expressing recombinant DENV-2 RUN-18 NS1 protein served as a control. In (**A**), the mean fluorescence intensity (MFI) of FITC signal in cells was examined by FACS analysis. For the detection of the E protein, cells were labeled using both mAbs 4G2 (left) and 4E11 (right). The results are the mean (± SE) of three independent assays in duplicate. Statistical analysis for prM-E was performed and noted (**** *p* < 0.0001). In (**B**), immunoblot assay on RIPA cell lysates were performed for detection of the E protein using mAb 4G2. Data are representative of three independent experiments.

The cytotoxicity of recombinant RUN-18 prM-E heterodimer was evaluated by quantifying lactate dehydrogenase (LDH) or MTT activity in A549 cells transfected with pcDNA3/RUN-18 prME for 18 h (Figure 2). Plasmid pcDNA3/RUN-18 NS1 was used as a control. A comparable level of LDH (Figure 2A) and MTT (Figure 2B) activity was found in A549 cells expressing prM and E proteins or NS1 protein. Thus, expression of recombinant RUN-18 envelope proteins in A549 cells transfected with pcDNA3/RUN-18 prME for 18 h is a suitable model system for studying the impact of amino-acid substitution M-(I36V) on the behavior of the E protein.

**Figure 2.**
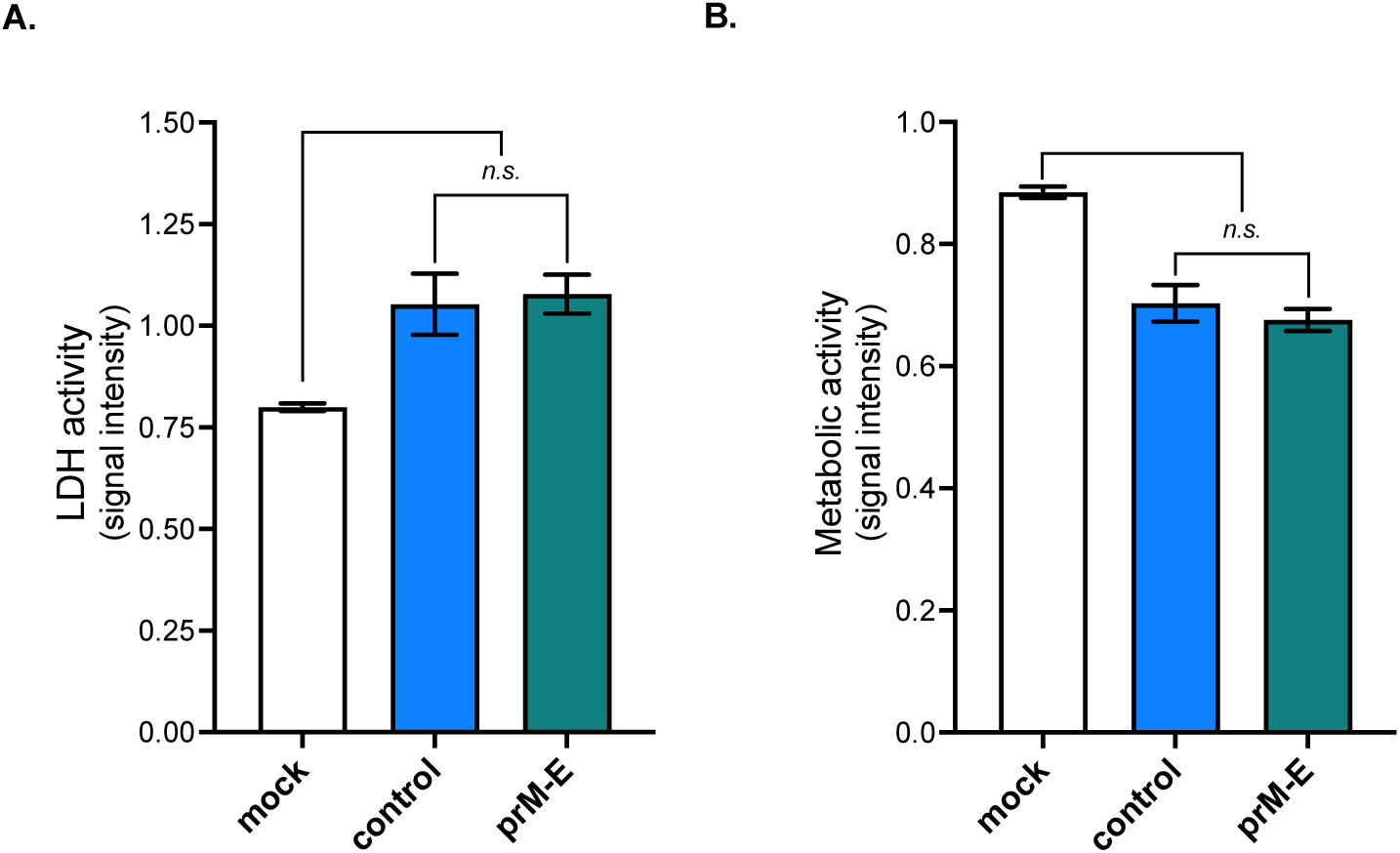
Cytotoxity of recombinant RUN-18 prM and E proteins. A549 cells were transfected for 18 h with plasmid pcDNA3/RUN-18 prME expressing recombinant RUN-18 prM and E proteins (prM-E) or mock-transfected (mock). A plasmid expressing recombinant DENV-2 RUN-18 NS1 protein served as a control. In (**A**), LDH activity was measured and O.D. values were expressed as signal intensity. In (**B**), MTT activity was measured and O.D. values were expressed as signal intensity. The results are the mean (± SE) of three independent assays with replicates. Statistical analysis for comparing mock, prM-E and NS1 was performed and noted (** *p* < 0.01, * *p* < 0.05; *n.s*: not significant).

### The M-(I36V) substitution influences RUN-18 E protein

To assess the impact of a Val residue at position M-36 on the expression of RUN-18 E protein, site-directed mutagenesis was performed on plasmid pcDNA3/RUN-18 prME to generate a mutant plasmid bearing Ile-to-Val substitution at position M-36 (Figure 3A). We also obtained two additional control plasmids bearing the single mutation M-(I36A) or the triple substitutions M-(E33A, W35A, R38A) (Figure 3A). It is worth noting that residues E33/W35/R38 are conserved among DENV-2 M proteins with a particular emphasis for W35 which is unchanged among flaviviruses [18]. We assessed whether structural information can predict the impact of amino-acid substitutions on RUN-18 M protein conformation. The M protein consists of a N-terminal ectodomain (residues 1/40) with a central α-helix followed by two transmembrane domains TMD-1 and TMD-2 which compose the C-terminal transmembrane region of the protein. The M residues 33 to 38 form the α-helix of the M ectodomain. Three-dimensional structure of RUN-18 M protein was performed by modeling on Phyre2 protein fold recognition server (Figure 3B). Modeling mutation in RUN-18 M protein suggests that structure is unchanged in M mutants bearing the triple substitutions M-(E33A, W35A, R38A) or single substitutions M-(I36A) and M-(I36V). Noteworthy *in silico* structure analysis showed that the different amino-acid substitutions in RUN-18 M ectodomain may have no impact on the α-helix conformation (Figure S2).

**Figure 3.**
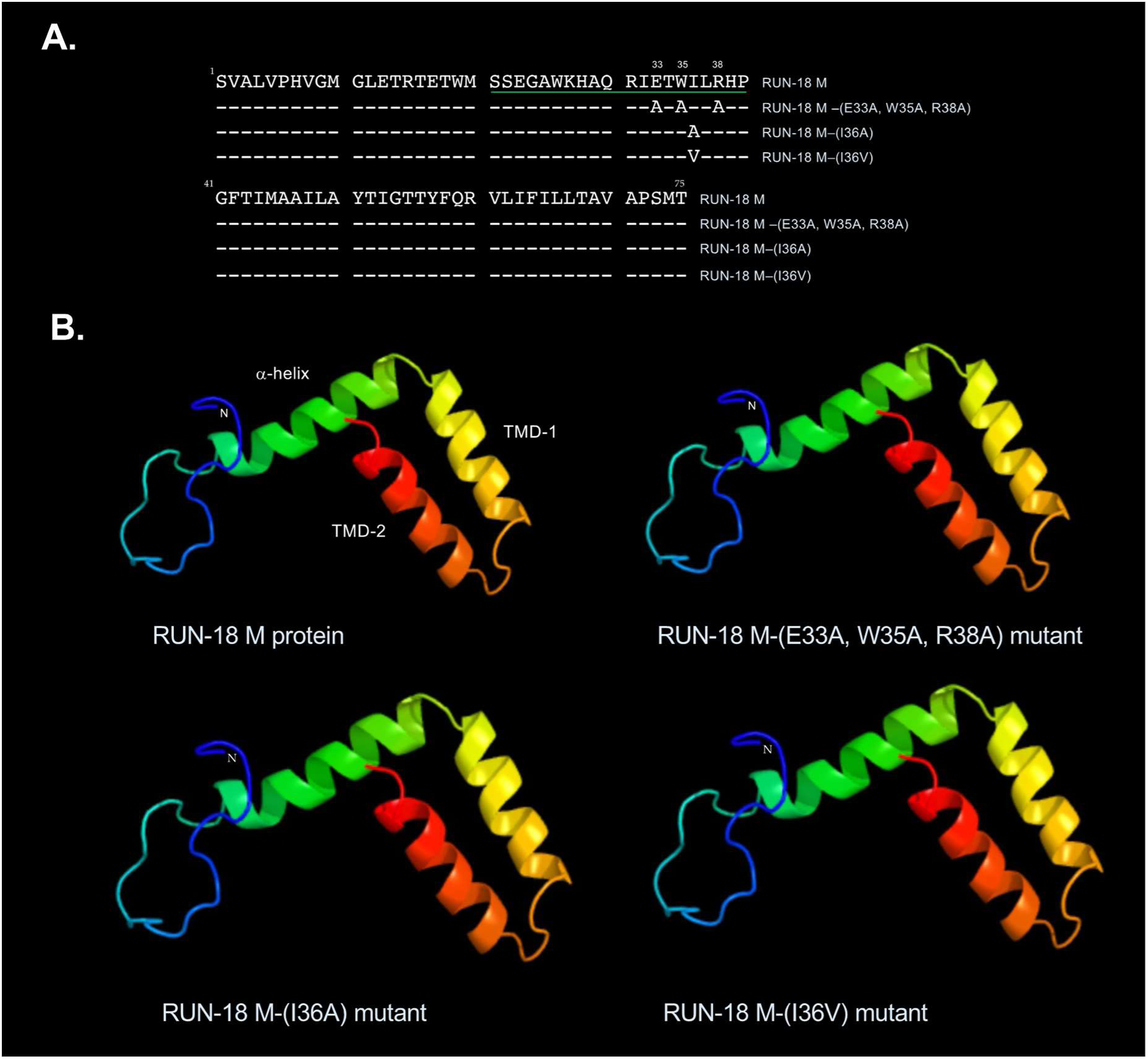
Impact of amino-acid substitutions on RUN-18 M structure. In (**A**), alignment of RUN-18 M protein (75 aminoacid residues) and mutants bearing the amino-acid substitutions at positions M-33, M-35, M-36 and M-38. The 20-residue peptide □-helix in M ectodomain is underlined in green. In (**B**), tridimensional structure prediction of RUN-18 M protein and the three mutants bearing the triple substitutions M-(E33A, W35A, R38A), the mutation M-(I36A), or the mutation M-(I36V) was performed by modeling on Phyre2 protein recognition server (http://www.sbg.bio.ic.ac.uk/~phyre2/html/page.cgi?id=index) based on template EMDB-5499 data. The predicted structures were analyzed using JSmol, a program for visualization of tridimensional molecules. The RUN-18 M protein consists of a N-terminal ectodomain (residues 1/40) with a central α-helix (green) followed by two transmembrane domains TMD-1 (yellow) and TMD-2 (red) which form the C-terminal transmembrane region of the protein.

Human epithelial A549 cells were transfected for 18 h with pcDNA3/RUN-18 prME or each of the three plasmid mutants. By RT-qPCR analysis on total RNA extracted from transfected cells, we checked that the different plasmids produced similar amounts of mRNA transcripts coding for prM and E proteins (Figure S3). FACS analysis showed that introduction of alanine residues at positions M-33/35/38 greatly reduced the reaction of anti-E mAbs 4G2 and 4E11 with RUN-18 E protein (Figure 4). Thus, amino-acid substitutions in RUN-18 ectoM might have a major impact on antigenic reactivity of E protein. The substitution of isoleucine by valine but not alanine at position M-36 greatly affected the reactivity of the two anti-DENV-2 E mAbs with RUN-18 E protein (Figure 4). This suggests that M residue 36 impact on the DENV-2 E protein could rely on the intrinsic nature of hydrophobic amino-acid at this position.

**Figure 4.**
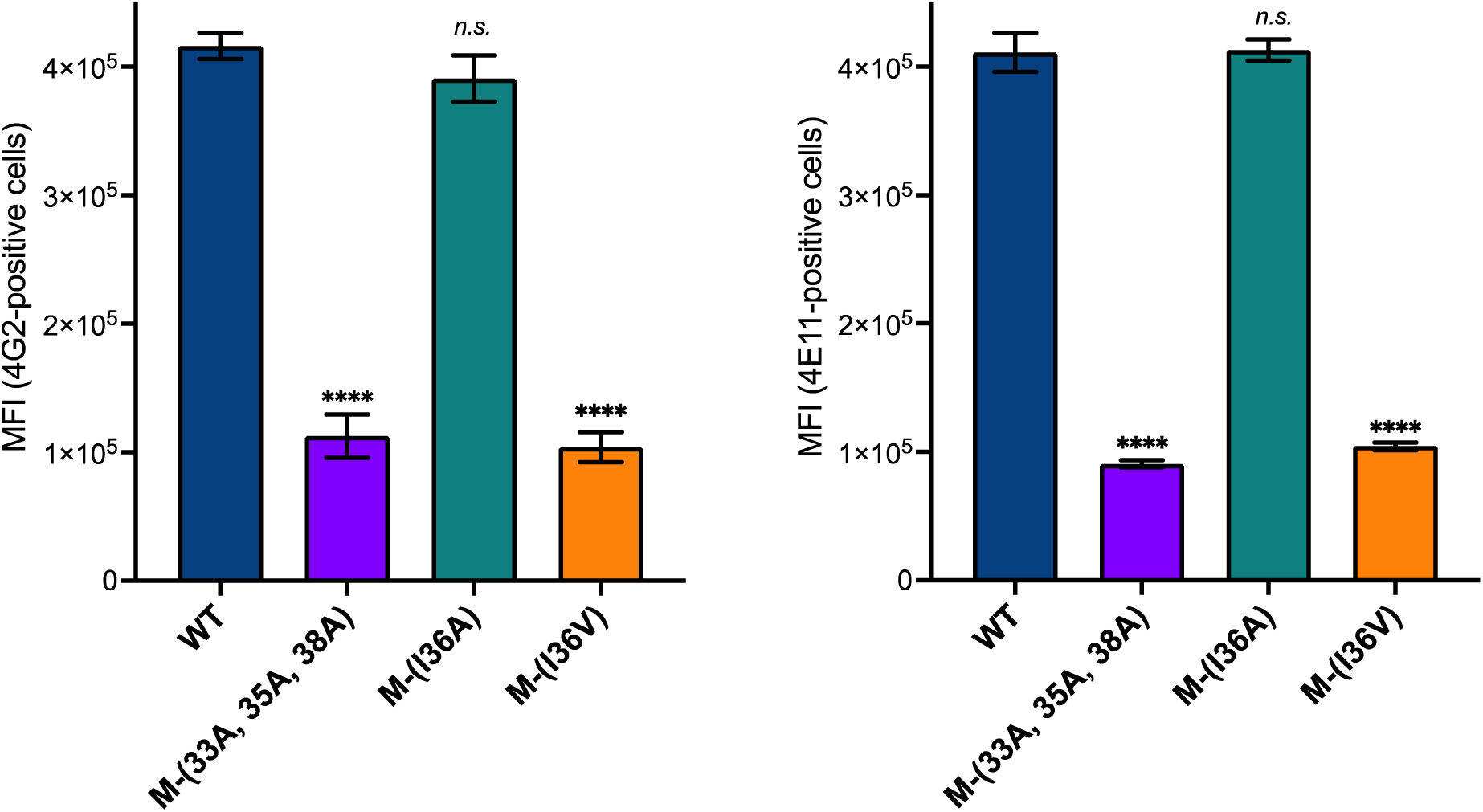
Impact of M amino-acid substitutions on antigenic reactivity of RUN-18 E protein. A549 cells were transfected 18 h with plasmid pcDNA3/RUN-18 prME expressing recombinant DENV-2 RUN-18 prM and E proteins (WT) or mutants bearing triple substitutions M-(E33A, W35A, R38A) or single substitutions M-(I36A) and M-(I36V). FACS analysis was performed on RUN-18 E protein using anti-DENV E mAb 4G2 (left) and 4E11 (right). Mean intensity fluorescence (MFI) of intracellular E protein was determined and the results are the mean (± SE) of three independent assays in duplicate. Statistical analysis for comparing WT and the mutants was performed and noted (**** *p* < 0.0001; *n.s*.: not significant).

As our data suggest that Ile-to-Val substitution at position M-36 affects RUN-18 E protein expression, we then examined whether this substitution could alter the subcellular distribution of the protein. Confocal immunofluorescence (IF) microscopy using mAb 4G2 was assessed on A549 cells expressing recombinant RUN-18 prM and E proteins (Figure 5). The accessibility of E protein into the endoplasmic reticulum (ER) compartment was evaluated by IF using ant-calnexin (CNX) antibody. There was a clear co-localization of RUN-18 E protein with ER-resident CNX (Figure 5). Dotted E protein-related foci which do not co-colocalize with CNX were also detected in A549 cells suggesting a trafficking of E protein through the secretory pathway. We noted that the triple substitutions M-(E33A, W35A, R38A) resulted in accumulation of RUN-18 E protein in the ER compartment undergoing an apparent morphology remodeling (Figure 5). There was also a lack of dotted E protein-related foci suggesting a retention of the protein in the ER. Introduction of valine but not alanine at position M-36 resulted in accumulation of E protein in the perinuclear region (Figure 5). Thus, the M-(I36V) mutation may drive important changes in the subcellular localization of RUN-18 E protein in A549 cells.

**Figure 5.**
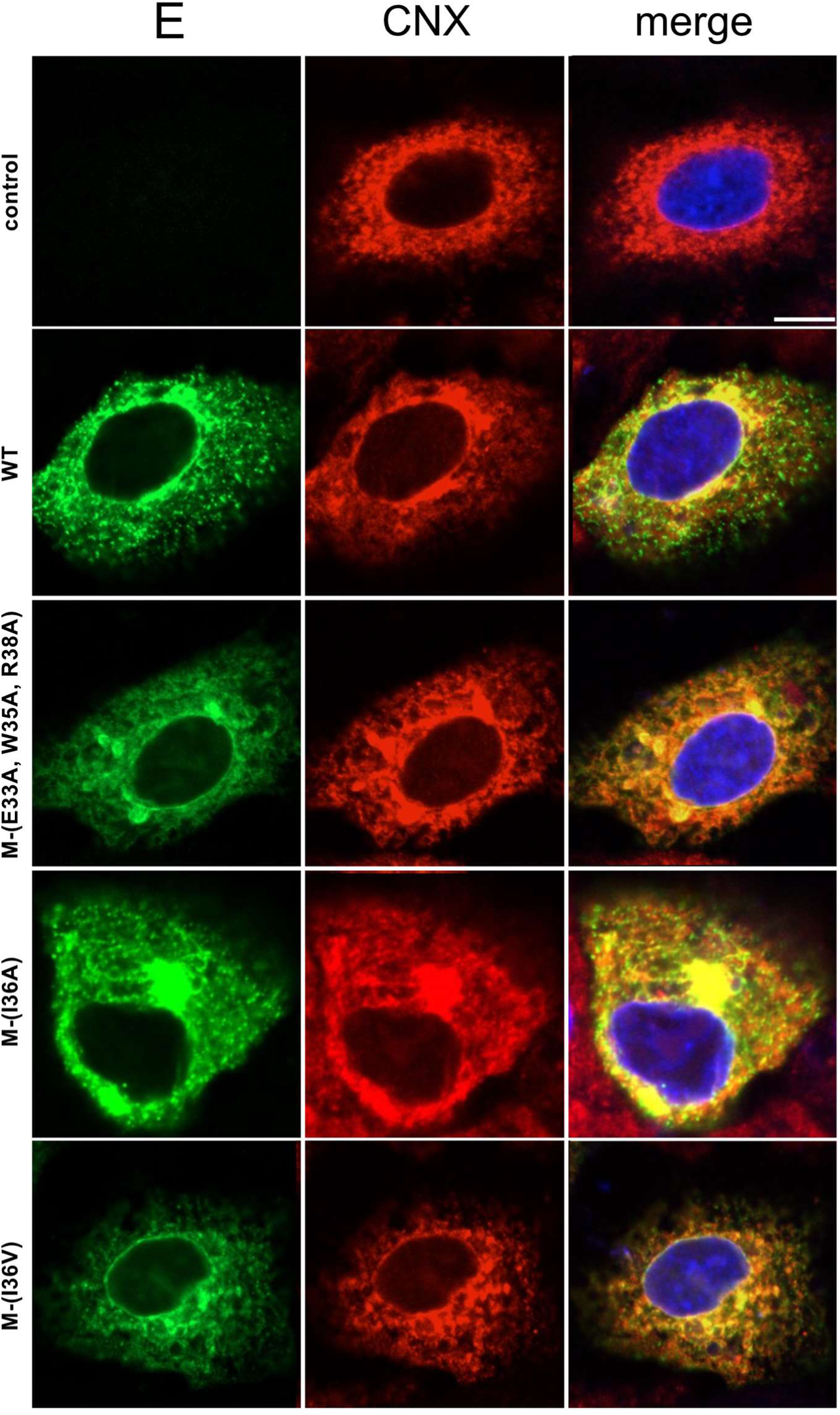
Subcellular distribution of RUN-18 E protein co-expressed with prM. A549 cells were transfected 18 h with plasmid pcDNA3/RUN-18 prME expressing recombinant DENV-2 RUN-18 prM and E proteins (WT) or mock-transfected (control). Effects of triple substitutions M-(E33A, W35A, R38A) and single mutation M-(I36A) or M-(I36V) on subcellular distribution of recombinant E protein were tested in A549 cells using mutant plasmids derived from pcDNA3/RUN-18 prME. For confocal microscopy analysis, cells were fixed, permeabilized and then immunolabeled using anti-DENV E mAb 4G2 and anti-calnexin (CNX) antibody. Cells were visualized for E (green) and CNX (red). Merged pictures are shown in the right panels. Scale bar, 5 μm. Data are representative of three independent experiments.

Considering that M-(I36V) mutation impacts the proper folding of RUN-18 E protein in A549 cells, we evaluated whether substitution also has an effect on viral proteins cytotoxicity (Figure 6). A549 cells were transfected for 30 h with plasmids expressing RUN-18 prM and E proteins bearing the triple substitutions M-(E33A, W35A, R38A), or the single substitutions M-(I36A) and M-(I36V). Measurement of LDH (Figure 6A) or MTT (Figure 6B) activity showed that introduction of valine but not alanine at position M-36 prevented the loss of cell viability associated with prM and E expression. A similar effect has been observed with the triple substitutions M-(E33A, W35A, R38A) (Figure 6). These results showed that Ile-to-Val substitution at position M-36 is sufficient to decrease the cytotoxicity of RUN-18 prM and E proteins in A549 cells.

**Figure 6.**
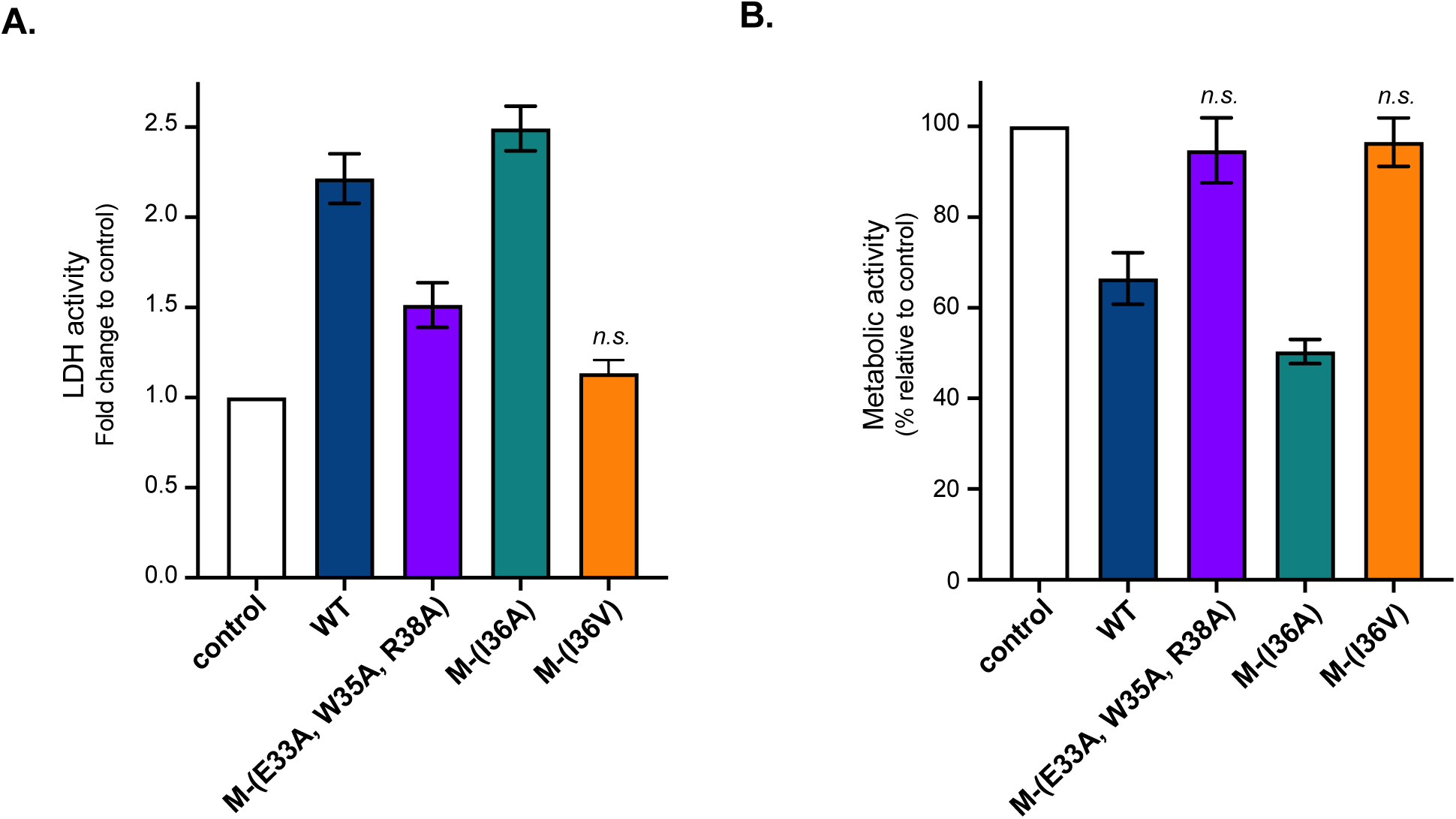
Impact of M amino-acid substitutions on the cytotoxicity of RUN-18 prM and E proteins. A549 cells were transfected with plasmid pcDNA3/RUN-18 prME expressing recombinant DENV-2 RUN-18 prM and E proteins (WT) or mock-transfected (control) for 30 h. Impact of single substitutions M-(I36A) and M-(I36V) and triple substitutions M-(E33A, W35A, R38A) on cytotoxicity of prM and E proteins was tested in A549 cells using mutant plasmids derived from pcDNA3/RUN-18 prME. In (**A**), LDH activity was measured and cell membrane permeability was expressed as a fold change relative to control. In (**B**), the level of cell metabolic activity was measured using MTT assay and expressed as percentage relative to control. The results are the mean (± SE) of three independent assays in four replicates. Statistical analysis for comparing control with WT and the three M mutants was performed and noted (**** *p* < 0.0001, *** *p* < 0.001, ** *p* < 0.01, * *p* < 0.05; *n.s*.: not significant).

We assessed whether expression of RUN-18 prM and E proteins induce apoptosis pathway in A549 cells. By measuring enzymatic activity of caspases-3 and −7 at 30 h post-transfection, we observed that expression of RUN-18 prM and E proteins can trigger apoptosis (Figure 7). The triple substitutions M- (E33A, W35A, R38A) significantly reduced the capacity of RUN-18 prM and E proteins to induce apoptosis. Remarkably, the Ile-to-Val substitution at position M-36 can abrogate the apoptotic effect of RUN-18 prM and E proteins in A549 cells (Figure 7). As expected, Ile-to-Ala substitution at position M-36 has no effect on the death-promoting activity of RUN-18 prM and E proteins. Taken together, these results suggest that Ile-to-Val substitution at position M-36 affect RUN-18 E protein expression associated with loss of cytotoxicity

**Figure 7.**
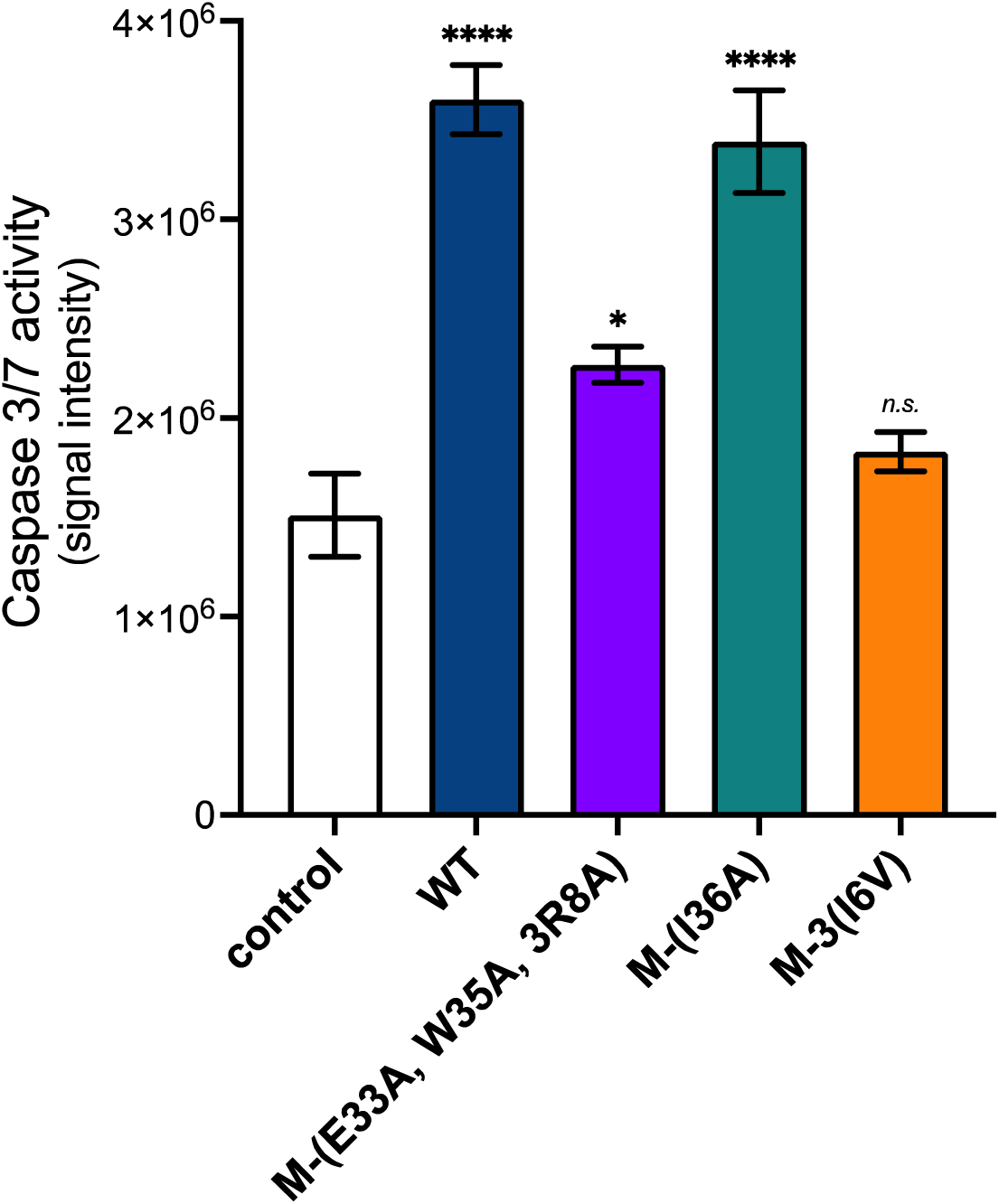
Impact of M amino-acid substitutions on apoptosis triggered by RUN-18 prM and E proteins. A549 cells were transfected with plasmid pcDNA3/RUN-18 prME expressing recombinant DENV-2 RUN-18 prM and E proteins (WT) or mock-transfected (control) for 30 h. Impact of single substitutions M-(I36A) and M-(I36V) and triple substitutions M- (E33A, W35A, R38A) on apoptosis triggered by prM and E proteins was tested using mutant plasmids derived from pcDNA3/RUN-18 prME. Caspase-3 and −7 enzymatic activity was measured and O.D. values were expressed as signal intensity. The results are the mean (± SE) of three independent assays in four replicates. Statistical analysis for comparing control with WT and the three M mutants was performed and noted (**** *p* < 0.0001, * *p* < 0.05; *n.s*.: not significant).

### The M-(I36V) mutation potentiates cell death-inducing activity of RUN-18 ApoptoM

The DENV-2 M residues 31 to 41 compose a pro-apoptotic peptide referred as ApoptoM [16]. We recently reported that RUN-18 ApoptoM linked to the C-terminus of secreted form of green fluorescent protein (sGFP) can trigger apoptosis in A549 cells [22]. Given that amino-acid substitutions on residue M-36 impact ApoptoM cytotoxicity [16], we evaluated the effect of Ile-to Val substitution on the apoptosis triggered by ApoptoM. Consequently, we generated the plasmid pcDNA3/sGFP-ApoptoM expressing RUN-18 ApoptoM linked to the C-terminus of GFP and preceded by the prM signal peptide (Figure 8). By directed mutagenesis, the single substitutions M-(I36V) and M-(I36A) or triple substitutions M-(E36A, W35A, R38A) were introduced in the sGFP-ApoptoM construct to generate ApoptoM-(I36V), ApoptoM- (I36A), and ApoptoM-(E33A, W35A, R38A) mutants, respectively (Figure 8). As a control, we used the sGFP construct lacking ApoptoM peptide at the C-terminus.

**Figure 8.**
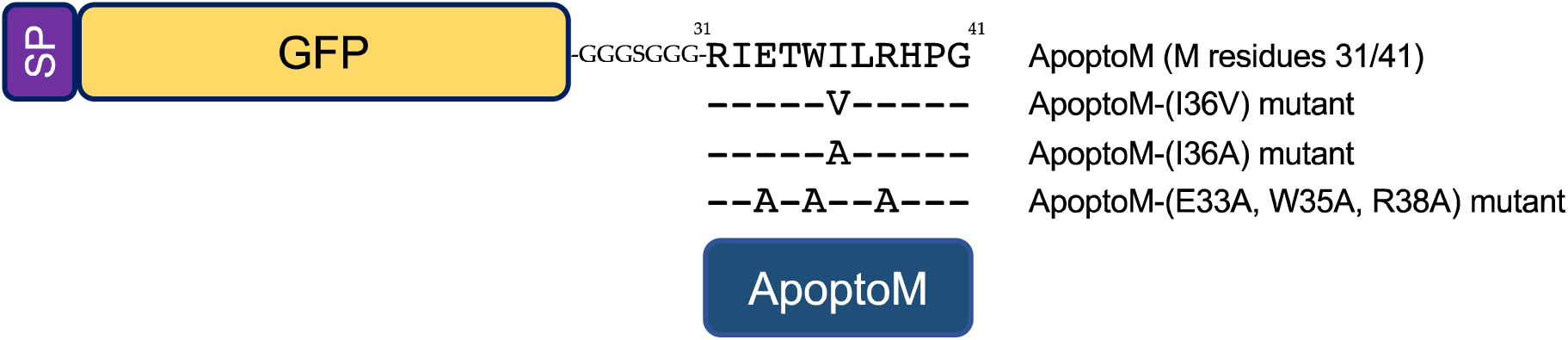
Schematic representation of sGFP-ApoptoM constructs. The DENV-2 RUN-18 M residues 31 to 41 referred as RUN-18 ApoptoM peptide are fused to the C-terminus of GFP reporter protein using a short Gly-Ser sequence spacer. The sGFP-ApoptoM construct is preceded by the RUN-18 prM signal peptide (SP). The sGFP-ApoptoM construct mutants were made by replacing the residues M-E33/W35/W38 with alanine or residue M-36I with valine or alanine. The different sGFP-ApoptoM constructs were inserted into plasmid pcDNA3.

We first determined the subcellular distribution of sGFP-ApoptoM expressed in A549 cells. In a first round of analysis, confocal IF microscopy was performed on A549 cells transfected with control plasmid expressing sGFP for 18 h (Figure 9A). A co-localization of sGFP with ER-resident calnexin (CNX) has been observed in A549 cells verifying that soluble reporter protein was correctly addressed to the ER compartment (Figure 9A). In A549 cells transfected with plasmid expressing sGFP-ApoptoM, large dots of GFP-related structures that partially co-localize with CNX were visualized in the ER which displays a coarse reticular network at the cell periphery (Figure 9B). The formation of these GFP-related structures might be due to sGFP-ApoptoM aggregation into the ER compartment leading to morphology changes. The replacement of RUN-18 ApoptoM by RUN-18 ectoM as sGFP-ectoM fusion protein resulted in the formation of a crown-like structure (CLS) accompanied by morphological changes in the ER (Figure 9C). Thus, the expression of the last ectoM residues which form ApoptoM causes rearrangement of the ER by favoring CLS formation.

**Figure 9.**
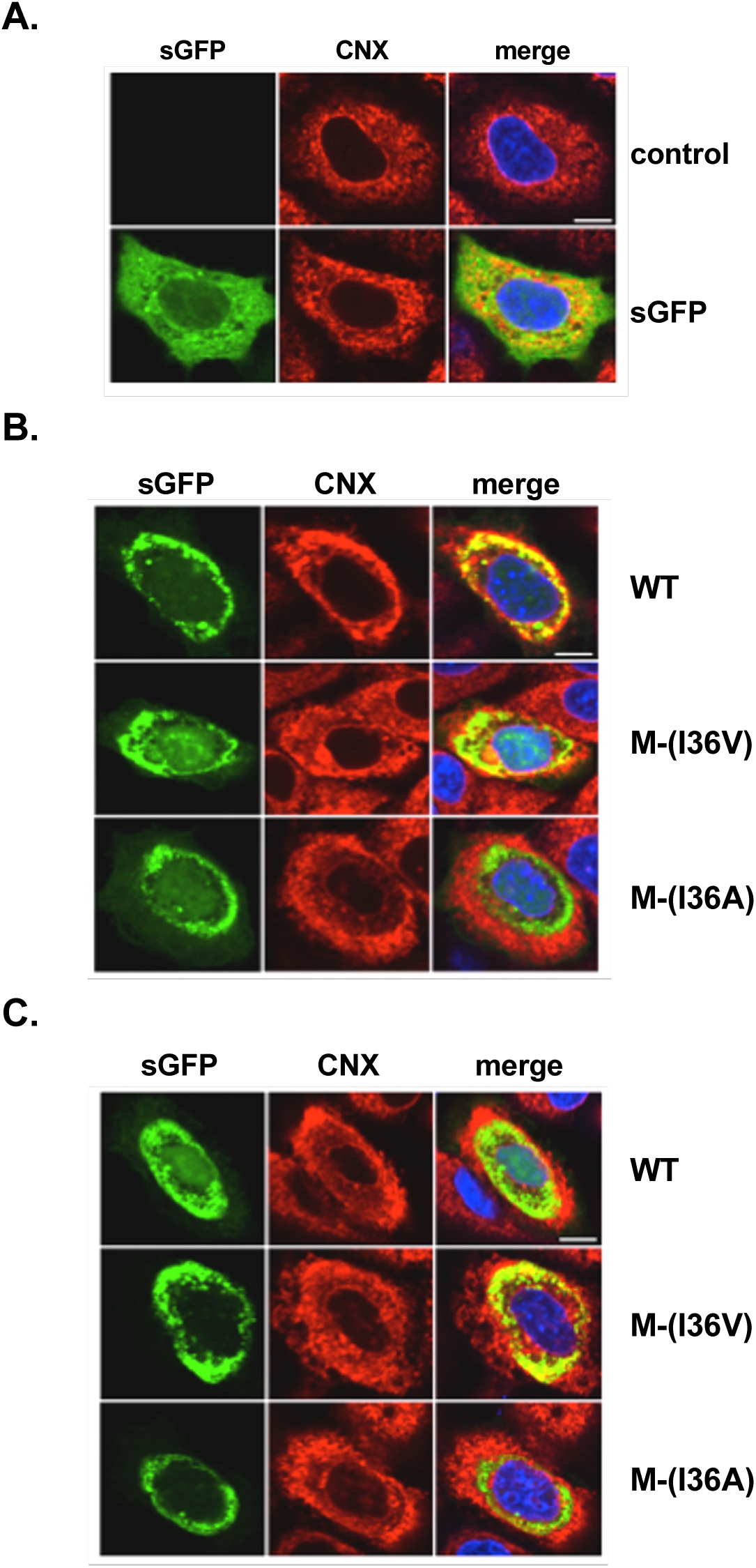
Subcellular distribution of sGFP-ApoptoM and sGFP-ectoM. A549 cells were transfected with plasmids expressing sGFP (**A**), sGFP-ApoptoM (**B**) or sGFP-ectoM (**C**) or mock-transfected (control) for 18 h. The ApoptoM (M residues 31-41) and ectoM (M residues 1-41) sequences are originated from DENV-2 RUN-18 strain. The isoleucine at position M-36 of ApoptoM or ectoM was replaced by valine or alanine. For confocal microscopy analysis, cells were fixed, permeabilized and then immunolabeled using anti-CNX antibody. Cells were visualized for soluble GFP (green) and CNX (red). Merged pictures are shown in the right. Scale bar, 5μm. Data are representative of three independent experiments.

We then addressed whether amino-acid substitutions at position M-36 affect the subcellular localization of sGFP-ApoptoM in A549 cells using mutant constructs carrying Ile-to-Val or Ile-to-Ala substitution at position M-36 (Figure 8). Introduction of M-(I36A) mutation in sGFP-ApoptoM resulted in a formation of a thinner CLS without alteration of ER morphology (Figure 9B). The attenuated phenotype for CLS was also observed in A549 cells expressing sGFP-ectoM mutant carrying Ile-to-Ala substitution at position M-36 (Figure 9C). There was a larger CLS and more severe ER rearrangement in A549 cells expressing sGFP-ApoptoM mutant carrying a valine at position M-36 compared to wildtype construct (Figure 9B). Effects of Ile-to-Val substitution at position M-36 on CLS formation and ER morphology were also observed with sGFP-ectoM mutant bearing the substitution (Figure 9C). Thus, RUN-18 ApoptoM bearing valine at position M-36 has a greater propensity to disrupt ER morphology compared to parental peptide.

To determine whether Ile-to-Val substitution at position M-36 impacts the cell death-inducing activity of DENV-2 ApoptoM, A549 cells were transfected with plasmids expressing sGFP-ApoptoM or mutants for 30 h (Figure 10). As controls, we used plasmids expressing sGFP-ApoptoM mutants bearing alanine at position M-36 or M-33/35/38 of viral peptide (Figure 8). Expression of wild-type sGFP-ApoptoM resulted in an increase in caspase-3/7 activity with a slight loss of cell viability (Figure 10A). Substitution of residues M-33/35/38 by alanine prevented ApoptoM-induced apoptosis (Figure 10A), leading to a loss of cytotoxicity (Figure 10B). Introduction of valine, and to a lesser extent alanine, at position M-36 resulted in a significant increase in caspase-3/7 activity (Figure 10A), and a decrease in cell metabolic activity reaching up 40% (Figure 10B). Thus, the substitution of isoleucine by valine at position M-36 potentiates the cell death-promoting capability of ApoptoM. Together, these results showed that RUN-18 ApoptoM is less efficient in inducing apoptosis compared with ApoptoM bearing valine or alanine at position M-36. The attenuated phenotype of RUN-18 ApoptoM was associated with a moderate effect on ER morphology.

**Figure 10.**
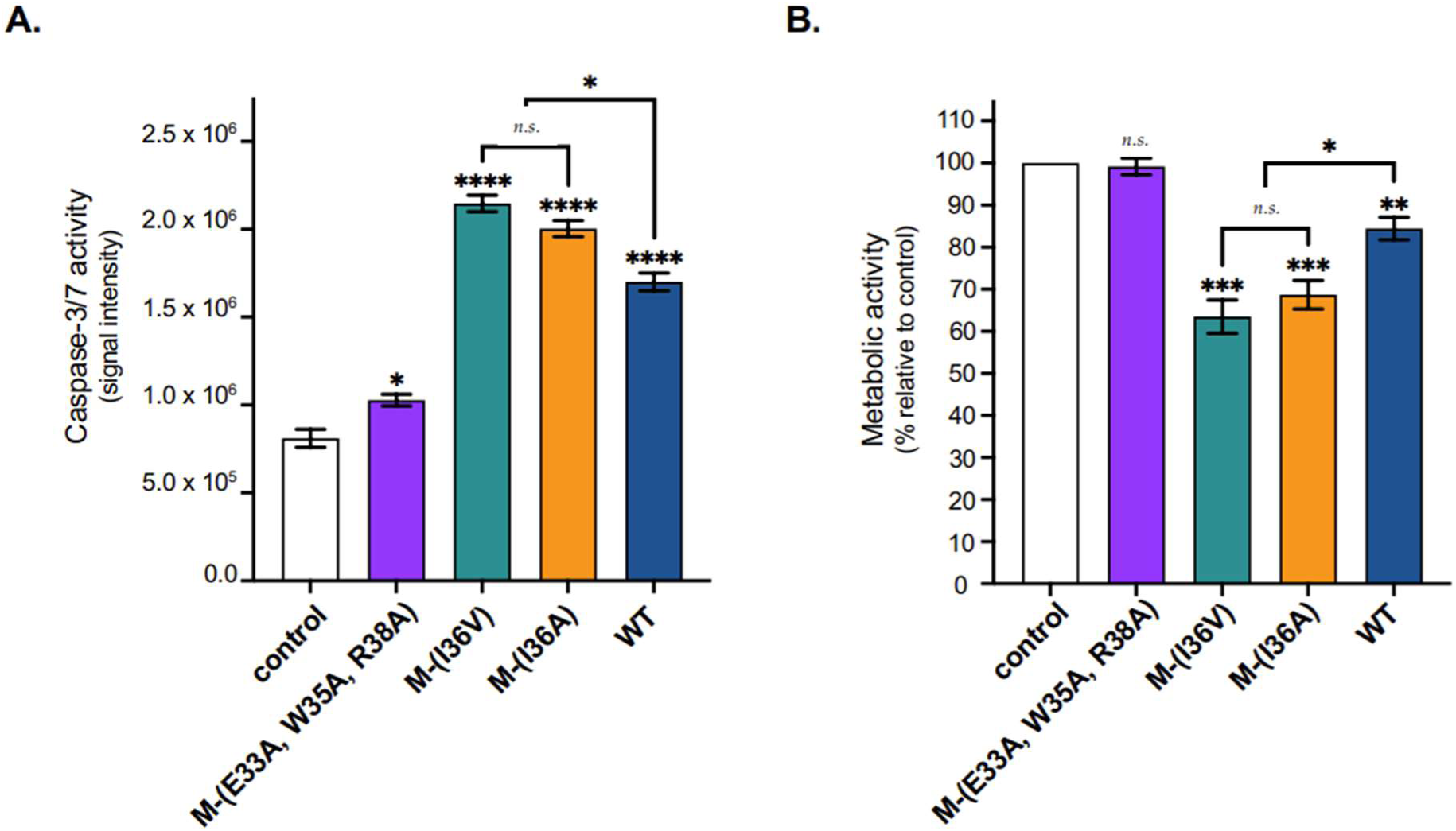
Impact of amino-acid substitutions on the cytoxicity of RUN-18 ApoptoM. A549 cells were transfected with plasmids expressing sGFP (control) or sGFP-ApoptoM (WT) and its mutants bearing triple substitutions M-(E33A, W35A, R38A) or single substitutions M-(I36A) and M-(I36V). In (**A**), caspase-3 and −7 enzymatic activity was measured and O.D. values were expressed as signal intensity. In (**B**), MTT assay was measured and expressed as percentage relative to control. The results are the mean (± SE) of three independent assays in four replicates. Statistical analysis for comparing control with WT and mutants was performed and noted (**** *p* < 0.0001, *** *p* < 0.001, ** *p* < 0.01, * *p* < 0.05; *n.s*.: not significant).

## DISCUSSION

Flavivirus prM/M protein (166 amino-acid residues) has been described to play important roles in various viral infection processes [23]. Although the mechanisms involved are not always well characterized, it is known that prM/M protein through its interaction with the E protein mediates assembly and maturation of viral particles [24,25]. The early stage of viral morphogenesis involves heterodimeric interactions between prM and E [26,27]. Throughout heterodimer prM-E, prM acts as a mandatory chaperone for proper folding of the E protein [28,29]. In the last stage of infectious life cycle, immature virus particles rely on prM cleavage at position 91 by furin in a post-Golgi compartment leading to “pr” peptide release and structural rearrangement in M-containing infectious extracellular particles [30]. Growing evidence suggests that mature M protein is involved in many aspects of flavivirus-induced pathogenesis. It is worth to note that expression of DENV M protein leads to inflammasome activation [31], while ZIKV M protein has been described to interact with cell plasma membrane acting as a viroporin to facilitate virus entry [32–34].

The surface membrane M protein is composed of an ectodomain (M residues 1/40) which forms an amphipathic peri-membrane α-helix (M residues 21/40) followed by two transmembrane domains. Inside the intracellular prM precursor protein, the M ectodomain plays an essential role in the heterodimeric interactions between prM and E [18,35–38]. It has been reported that glutamic acid at position M-33 is involved in prM/M interaction with Domain II of E protein [39]. Residues M-33/35/38 are highly conserved among flavivirus M proteins, with tryptophan at position M-35 being invariable [18]. Similarly, introduction of alanine at positions M-33/35/38 causes a severe defect in E protein folding. Previous reports showed that the nature of M residue 36 may have a major impact on flavivirus progeny production in mammalian cells [19,20,37]. The residue at position M-36 was essentially identified as an uncharged amino-acid with isoleucine (DENV-2, DENV-4, JEV, and WNV), alanine (DENV-1 and DENV-3), leucine (YFV) or valine (TBEV) [18], emphasizing that aliphatic hydrophobic properties of amino-acid residue in position 36 of M protein may have an impact on E protein expression.

Majority of DENV-2 M proteins have an Ile residue at position M-36 whereas only very few of them have Val residue [18]. Comparative sequence analysis between contemporary SWIO DENV-2 strains identified isoleucine at position M-36 for RUN-18 but valine for DES-14. This finding prompted us to evaluate whether Ile-to-Val substitution at position M-36 affects the proper folding of E protein. We first showed that substitution of valine at position M-36 altered the antigenic reactivity of RUN-18 E protein suggesting a major defect in heterodimeric interactions between prM and E proteins. Considering that substitution by an alanine has no effect on E protein, it is worth to think that intrinsic valine properties are involved in the observed defect of RUN-18 E protein expression.

We noted that DENV-2 RUN-18 and DES-14 E proteins differ by four amino-acid substitutions with a particular emphasis towards Gln-to-His substitution at position E-52 in the EDI/EDII interface and Ala-to-Thr substitution at position E-262 in EDII domain which faces the M ectodomain in the prM-E heterodimer (Table S1). Whether residues E-262T and E-52H could act as two compensatory amino-acid residues to the presence of valine at position M-36 in DES-14 is an important issue that remains to be investigated. Another important finding of the study was the lack of apoptosis induction in A549 cells expressing RUN-18 prM and E proteins with M protein bearing valine at position 36. Such a result was unexpected knowing that improperly folded viral envelope proteins usually give rise to severe ER dysfunction [40–42]. One hypothesis is that introduction of valine at position M-36 destabilizes the heterodimeric interactions between RUN-18 prM and E leading to a rapid proteolytic degradation of viral proteins.

RUN-18 M ectodomain includes a stretch of eleven amino-acids ^31^RIETWILRHPG^41^ with forms a pro-apoptotic viral peptide initially referred to as ApoptoM [16,17]. Translocation of ApoptoM into the ER and its subsequent transport through the secretory pathway is a pre-requisite for apoptosis induction [16]. In this study, we showed that substitution of alanine at positions M-33/35/38 abrogated the RUN-18 ApoptoM death-promoting activity. The M residue 36 acts on the capacity of ApoptoM to trigger apoptosis [16]. DENV-2 RUN-18 and DES-14 ApoptoM only differ at the position M-36 with Ile or Val residue, respectively. Using secreted GFP as reporter protein, different patterns of subcellular distribution of sGFP-ApoptoM protein have been identified in A549 cells according to the nature of M residue 36. Expression of RUN-18 ApoptoM linked to sGFP resulted in CLS formation associated with changes in ER morphology. It is thought that CLS was a consequence of interactions between ApoptoM and cytoskeletal motor protein dynein [43–45], which has been described as playing a role in trafficking of immature particles through its interaction with DENV-2 M protein [46]. Ile-to-Val substitution at position M-36 in RUN-18 ApoptoM coincided with the formation of larger CST and greater alteration of ER morphology. The substitution of isoleucine by valine, and a lesser extent alanine, at position M-36 leads to increase apoptosis-inducing activity of ApoptoM. Thus, DENV-2 ApoptoM is influenced by the hydrophobic nature of amino-acid residue in position M-36 most probably through protein-protein interactions capable of activating mitochondrial apoptosis pathway although mechanisms remain sill poorly understood [42].

Our data highlighted a major role for M residue 36 in proper folding of DENV-2 E protein and pro-apoptotic activity of ApoptoM. There is mounting evidence that small integral M protein is involved in flavivirus morphogenesis and infection processes [19,20,31,33,38]. It is therefore of priority to generate infectious molecular clones derived from RUN-18 and DES-14 to better understand the role of M residue 36 on the pathogenicity of contemporary SWIO DENV-2 strains. This would provide new information of interest for the development of future live-attenuated DENV vaccines.

## Supporting information

supplemental informations

## Author Contributions

J.D., G.G. and P.D. conceived and designed the experiments; JD performed the experiments; J.D., G.G. and P.D. analyzed the data; P.D. contributed to reagents/materials/analysis tools; J.D., G.G. and P.D. wrote the paper. All authors have read and agreed to the published version of the manuscript.

## Data Availability Statement

The data presented in this study are available in this article and Supplementary Materials.

## Acknowledgments

We thank P. Mavingui and C. El-Kalamouni for their interest in the study. We thank the members of MOCA team for helpful discussions.

