## supplemental informations for "The hydrophobic nature of dengue 2 M residue 36 influences E protein and ApoptoM peptide"

**Supplementary data**

**Table S1. Amino-acid substitutions between DENV-2 RUN-18 and DES-14 prM and E proteins.**

| Position | Antigenic domain | RUN-18 | DES-14 |
|---|---|---|---|
| prM-29 | n.a. | Asp | Asn |
| prM-127 (M-36) | n.a. | Ile | Val |
| E-52 | EDI/II junction | Gln | His |
| E-141 | EDI | Val | Ile |
| E-262 | EDII | Ala | Thr |
| E-322 | EDIII | Val | Ile |

n.a.: non applicable

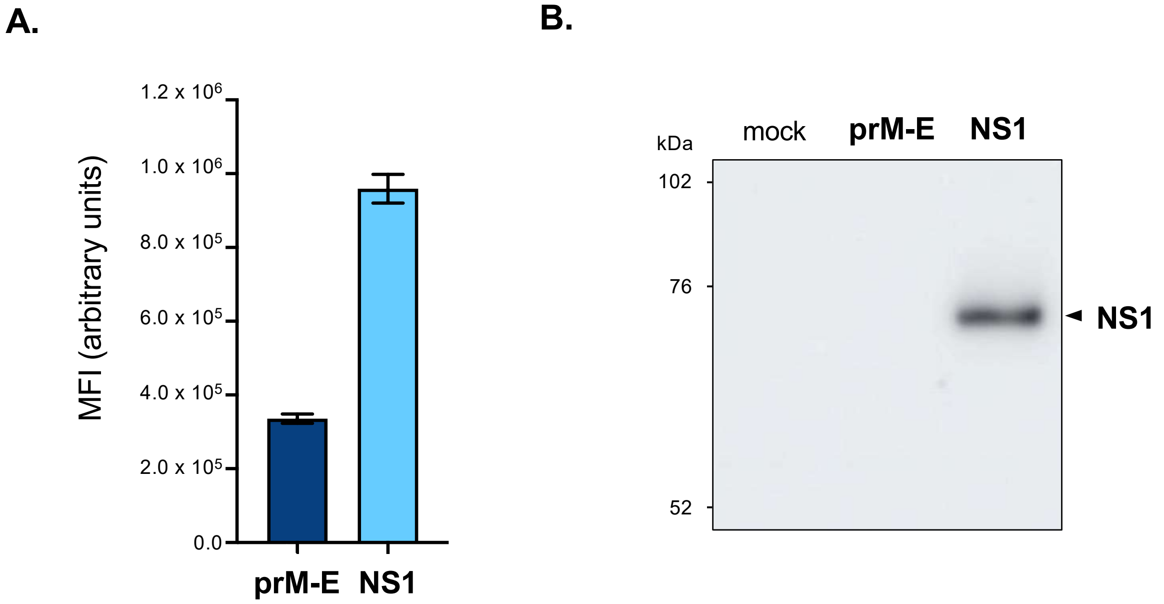


Figure S1. **Expression of RUN-18 NS1 protein in A549 cells.** A549 cells were transfected for 18 h with recombinant plasmids expressing recombinant RUN-18 prM and E proteins or NS1 protein, or mock-transfected (mock). In (**A**), for detection of the recombinant 6x(His)-tagged NS1 protein, transfected cells were labeled using anti-6x(His) mAb as primary antibody. MFI of FITC signal in cells was examined by FACS analysis. The results are the mean (± SE) of three independent assays. Statistical analysis for NS1 was performed and noted (*** $p < 0.001$). In (**B**), immunoblot assay on RIPA cell lysates were performed for detection of the NS1 protein using anti-6x(His) mAb as primary antibody.

**A.**

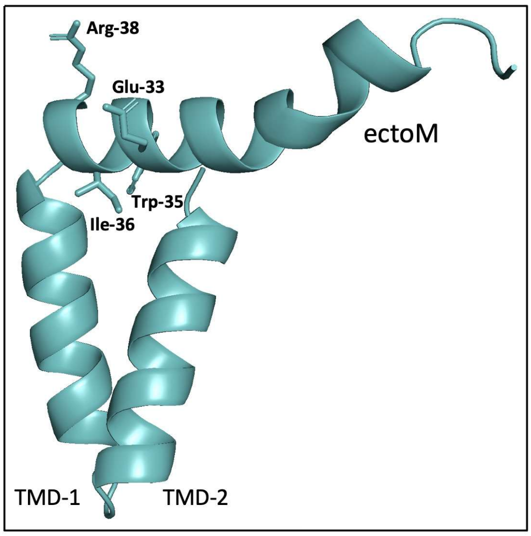


**B.**

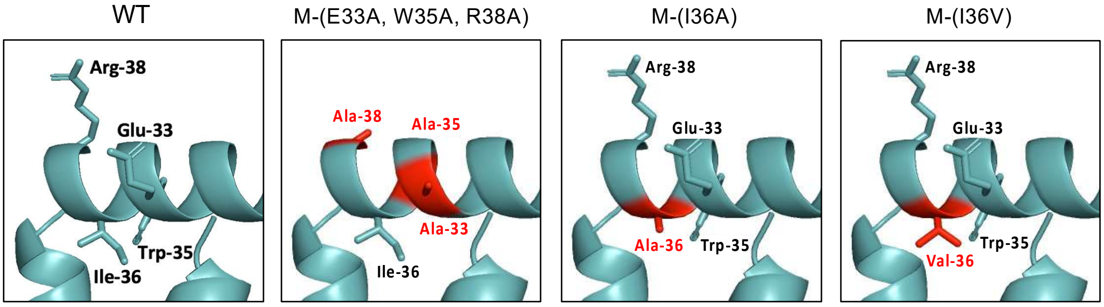


<u>Figure S2.</u> **Impact of amino-acid substitutions on $\alpha$-helix conformation of RUN-18 ectoM.** In (**A**), RUN-18 M protein structure was obtained from protein database bank (GenBank accession number MN272404.1) and rendered using three-dimensional structure from PDB (7KV8). All pictures were drawn using PyMol software. Amino-acid residues at the position M-33/35/36/38 are shown with sticks. In (**B**), a focus on $\alpha$-helix (M residues 21/40) of the RUN-18 ectoM (WT) showing the triple substitutions (M-E33A, W35A, R38A), and single substitutions M-(I36A) and M-(I36V).

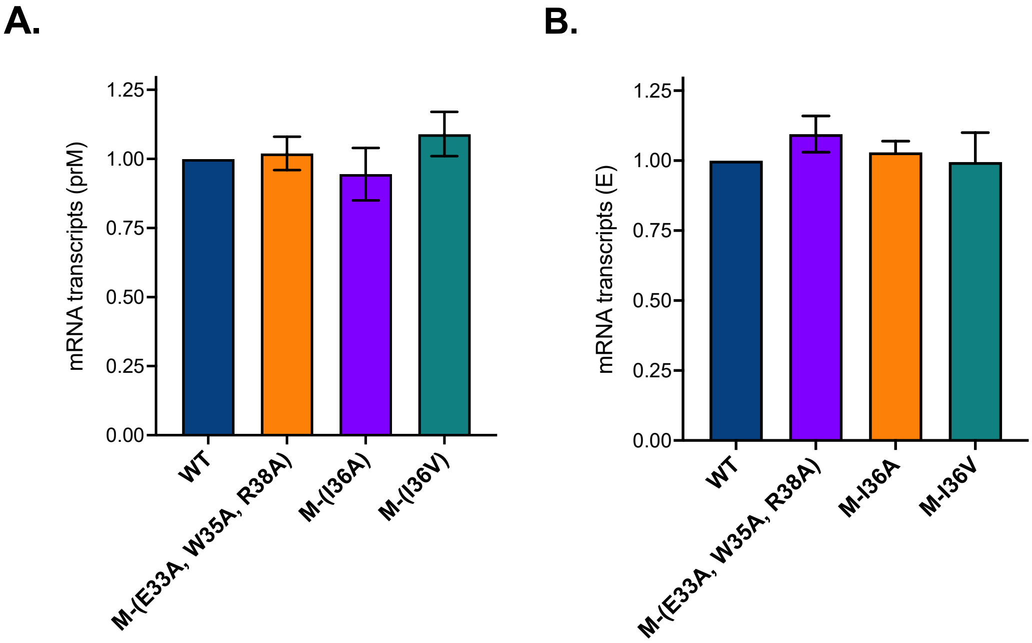


Figure S3. **RT-qPCR analysis on mRNA coding for recombinant RUN-18 prM and E proteins.** A549 cells were transfected 18 h with plasmid pcDNA3/RUN-18 prME (WT) or its three mutants bearing the triple substitutions M-(E33A, W35A, R38A), or the single substitutions M-(I36A) and M-(I36V). RT-qPCR assays were performed on total RNA extracted from transfected cells using specific primers for prM (**A**) or E (**B**) gene sequence. The results in ΔΔ*C*T values are the mean (± SE) of two independent assays. Statistical analysis for comparing mRNA copies encoding RUN-18 prM and E with the three M mutants was performed and differences that were not statistically significant are omitted.
